# PepGo: a deep learning and tree search-based model for *de novo* peptide sequencing

**DOI:** 10.1101/2025.02.24.640018

**Authors:** Yuqi Chang, Siqi Liu, Karsten Kristiansen

## Abstract

Identifying peptide sequences from tandem mass spectra is a fundamental problem in proteomics. Unlike search-based methods that rely on matching spectra to databases, *de novo* peptide sequencing determines peptides directly from mass spectra without any prior information. However, the design of models and algorithms for *de novo* peptide sequencing remains a challenge. Many *de novo* approaches leverage deep learning but primarily focus on the architecture of neural networks, paying less attention to search algorithms. We introduce PepGo, a *de novo* peptide sequencing model that integrates Transformer neural networks with Monte Carlo Tree Search (MCTS). PepGo predicts peptide sequences directly from mass spectra without databases, even without prior training. We show that PepGo surpasses existing methods, achieving state-of-the-art performance. To our knowledge, this is the first approach to combine deep learning with MCTS for *de novo* peptide sequencing, offering a powerful and adaptable solution for peptide identification in proteomics research.

## 1. Introduction

Mass spectrometry originated in the field of physics ^1^ but has since become widely used across numerous disciplines, including chemistry, biology, and medicine^2, 3^. In biology, mass spectrometry plays a crucial role in the analysis and identification of biomolecules such as proteins, lipids, and metabolites. For protein identification, the process begins with the extraction of proteins from a biological sample, followed by digestion into peptides using proteolytic enzymes (e.g., trypsin). These peptides are then charged and fragmented using techniques like collision-induced dissociation (CID), producing ions that generate tandem mass (MS/MS) spectra. From these spectra, the amino acid sequences of the peptides are identified. Peptide identification approaches can be broadly categorized into two main strategies based on their underlying principles: search-based approaches and *de novo* peptide sequencing.

Search-based approaches identify peptides by matching experimental MS/MS spectra to either theoretical spectra generated from a sequence database or real spectra from a spectral library. These methods rely heavily on a comprehensive database or library as a prerequisite reference^4^. In contrast, de novo peptide sequencing infers peptide sequences directly from MS/MS spectra without requiring a sequence database or spectral library.

Although search-based approaches dominate peptide identification in mass spectrometry-based proteomics, *de novo* peptide sequencing plays a critical role in identifying novel peptides or sequences absent from existing databases. For example, spectra that cannot be assigned a peptide sequence through searching have a chance to be rescued using *de novo* peptide sequencing. More importantly, due to the lack of corresponding sequences in the database or library, search-based approaches may underperform when analyzing peptides that have not been previously investigated, such as peptides from unsequenced organisms, or antigenic peptides presented by major histocompatibility complex(MHC) proteins.^5, 6^. In such cases, *de novo* peptide sequencing may be the only viable option for identification.

Table 1 summarizes the various approaches proposed for *de novo* peptide sequencing, categorized by their algorithms. Among these approaches, dynamic programming and graph theory-based algorithms are widely adopted. However, the inherent complexities of spectral data present challenges that simple models struggle to address. With the rapid development of high-performance hardware (such as GPUs) and advancements in methodology, artificial intelligence (AI) techniques have been applied to *de novo* peptide sequencing, showing promising results in recent experimental findings ^6–10^. Therefore, exploring the potential of AI techniques for tackling *de novo* peptide sequencing problems is highly valuable.

**Table 1:**
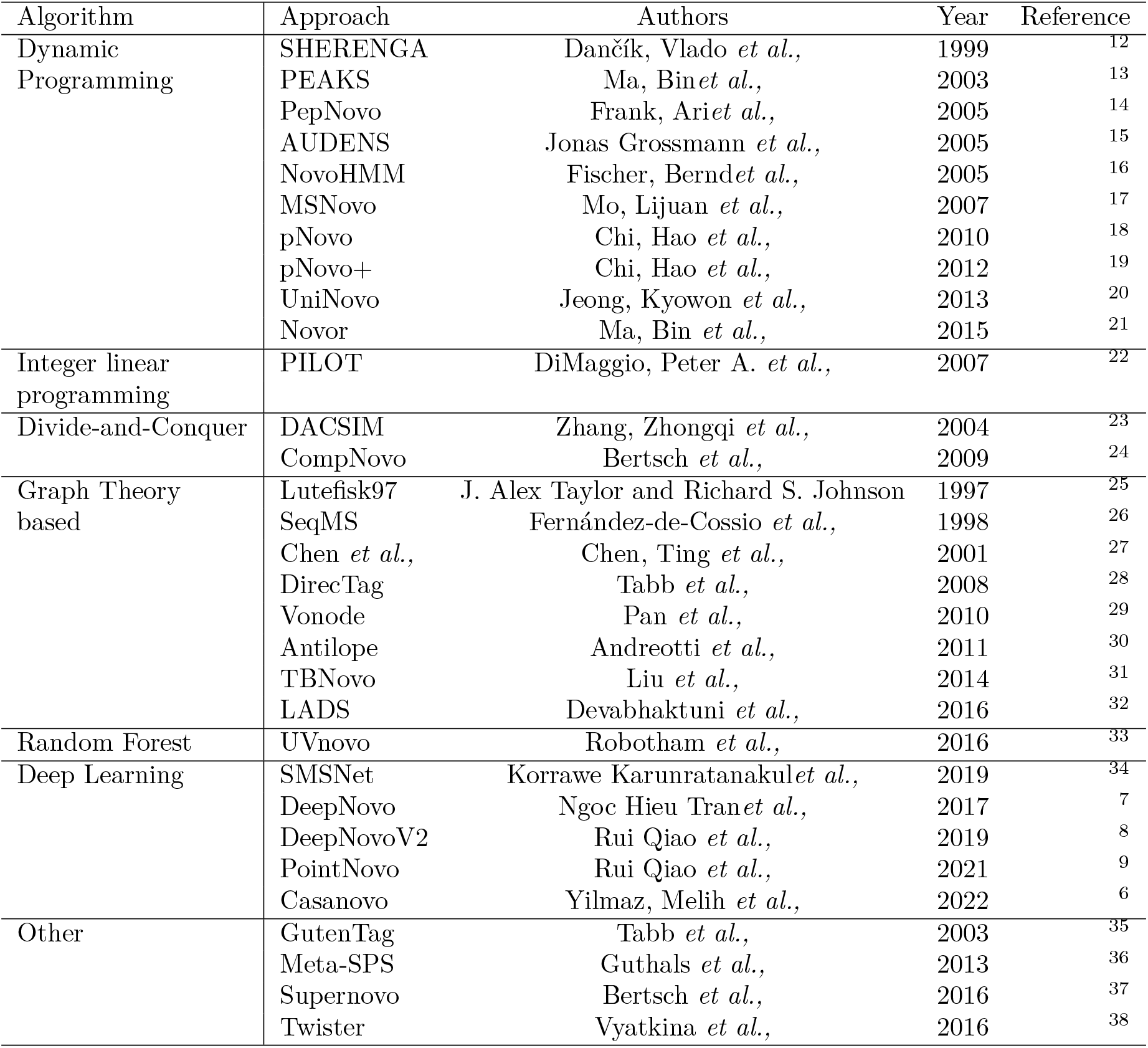
Summary of proposed *de novo* peptide sequencing approaches.

We introduce PepGo, a novel *de novo* peptide sequencing model that integrates Transformer neural networks with Monte Carlo Tree Search (MCTS). Inspired by AlphaGo’s groundbreaking combination of deep neural networks and tree search^11^, PepGo harnesses advanced AI techniques to achieve state-of-the-art accuracy in peptide sequencing. In the following sections, we systematically detail PepGo’s model architecture and algorithm design, evaluate its performance on benchmark synthetic peptide datasets and compare it against existing models to demonstrate its superior capabilities.

## 2. Methods

### 2.1 Peptide Representation

To rigorously represent peptides in a format suitable for computational processing, amino acid residues are denoted by their one-letter codes, with the symbol *<* representing the N-terminal and the symbol *>* representing the C-terminal. In this context, *<* and *>* correspond to the hydrogen (*H*) and hydroxyl (*OH*) groups at the two ends of the peptide, respectively. A residue with its post-translational modification (PTM) as a whole is denoted as the one-letter code plus the PTM (e.g., M+Oxidation). For simplicity, the term “*residue*” represents both unmodified residues and those with PTMs.

Conventionally, peptide sequences are presented in their natural, original order, from the N-terminal to the C-terminal. We refer to these as **forward peptide sequences**. Conversely, peptide sequences presented in the reverse order, from the C-terminal to the N-terminal, are called **reversed peptide sequences**.

Taking all these definitions into account, the forward peptide sequence GMRFWYK is represented as:

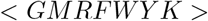

The reversed version of the same peptide sequence is:

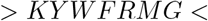

### 2.2. Spectrum Representation

The input to our model consists of mascot generic format (MGF) files, which contain multiple MS/MS spectra. Each spectrum is made up of a set of mass-to-charge ratio (*m*/*z*) and intensity (*I*) pairs that represent the fragment ions produced after collision, typically visualized as **peaks**. In a spectrum, peak *i* is represented by (*m*/*z*_*i*_, *I*_*i*_) and a spectrum containing *N* peaks is represented by 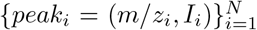. Figure 1 illustrates a theoretical spectrum where the peaks correspond to fragment ions from the peptide GMRFWYK. The data for this plot was simulated using MS2PIP^39^ and manually adjusted, making it an abstract example rather than being derived from any actual peptide.

**Figure 1.**
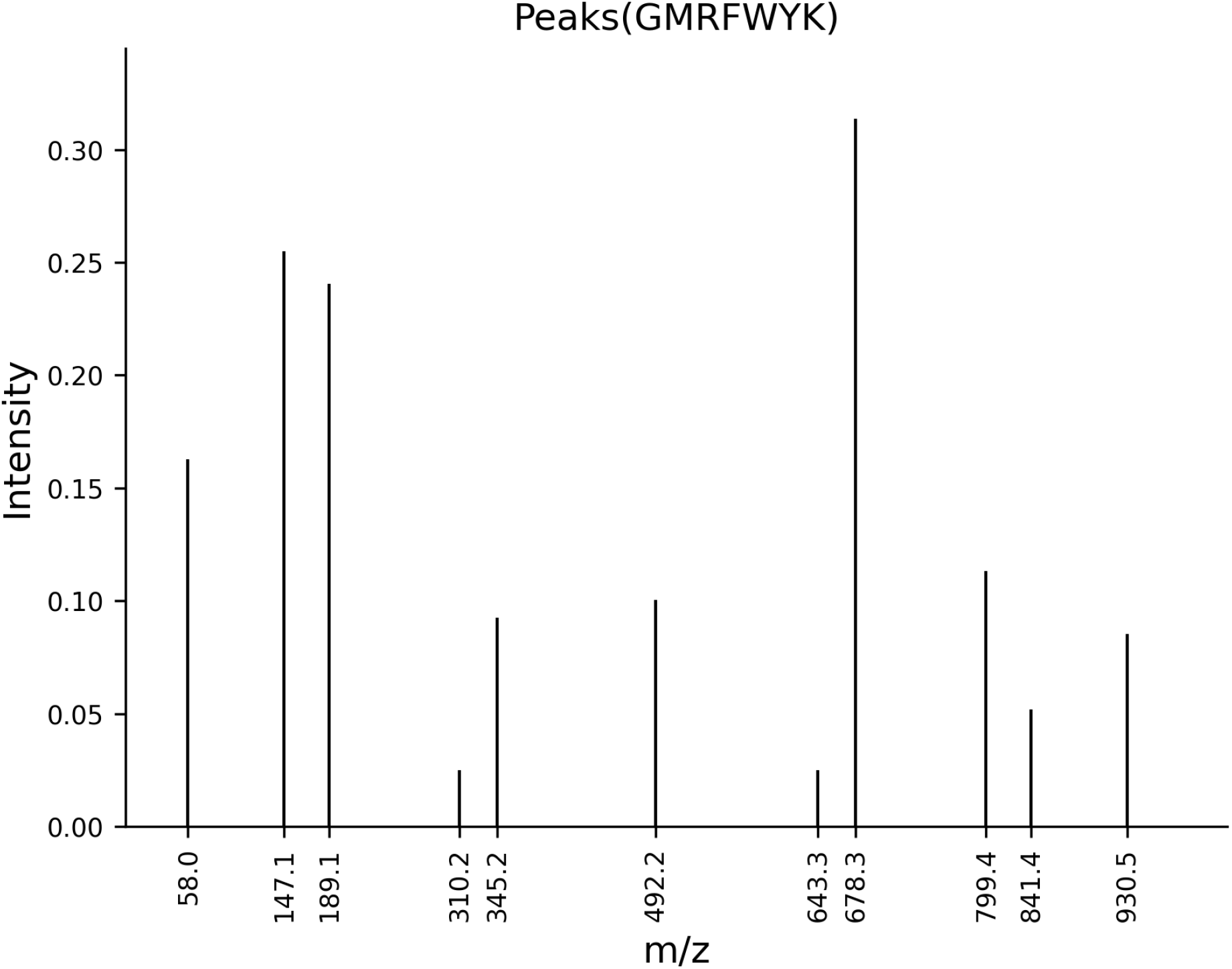
Theoretical spectrum delineating peaks corresponding to fragment ions from the peptide GMRFWYK

### 2.3 Probe Model

We define **spectrum probes** as the *m*/*z* values of peptide subsequences generated *in silico* following simulated ionization. The generation of spectrum probes involves two steps: first, a peptide subsequence is constructed by concatenating residues with the symbols *<* or *>*. Second, the constructed subsequence is ionized, taking into account different ion types, neutral losses, charges and other factors that influence the *m*/*z* values.

An example of probes generated from the subsequence <*G* is shown in Figure 2a. Initially, we assume the first residue in the N-terminal is *G*(glycine), so the subsequence <*G* is constructed. The subsequence <*G* is then ionized by adding one or two charges, resulting in dissociation into a,b, and y ions, without considering neutral losses. This process yields a set of six probes for subsequence <*G*. Table 2 provides a list of factors considered during simulated ionization.

**Table 2:**
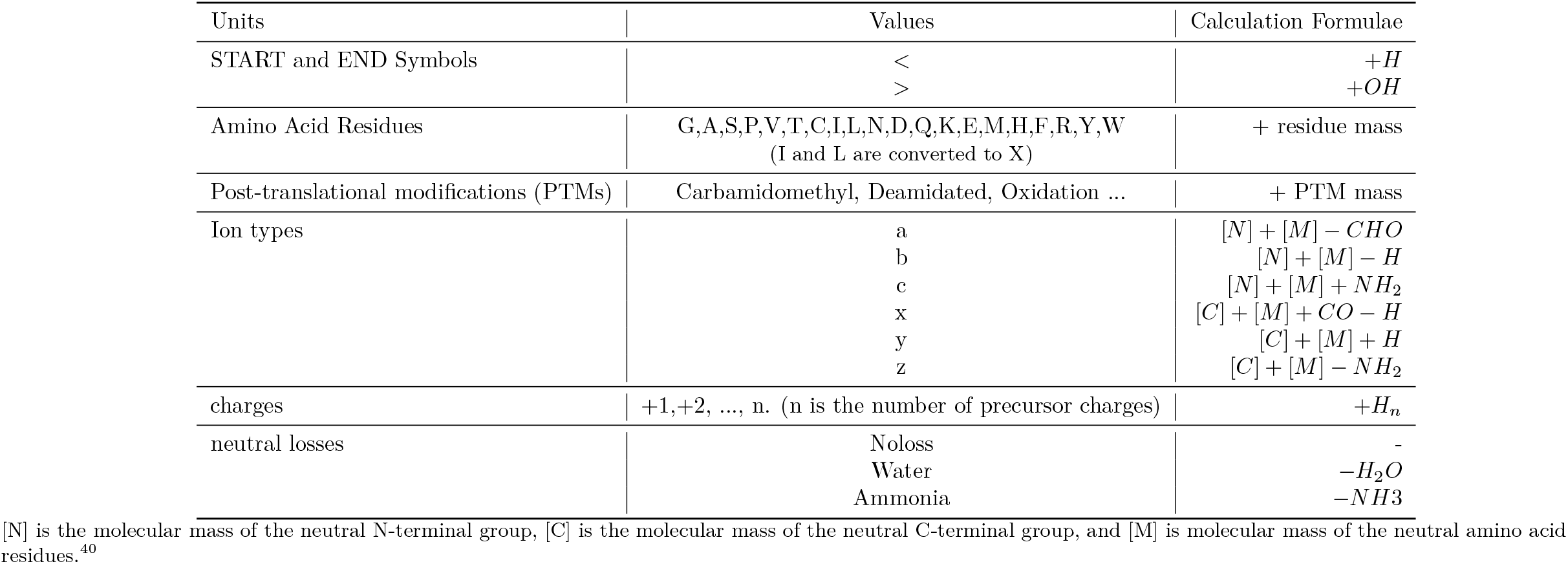
Factors considered in simulated ionization.

**Figure 2.**
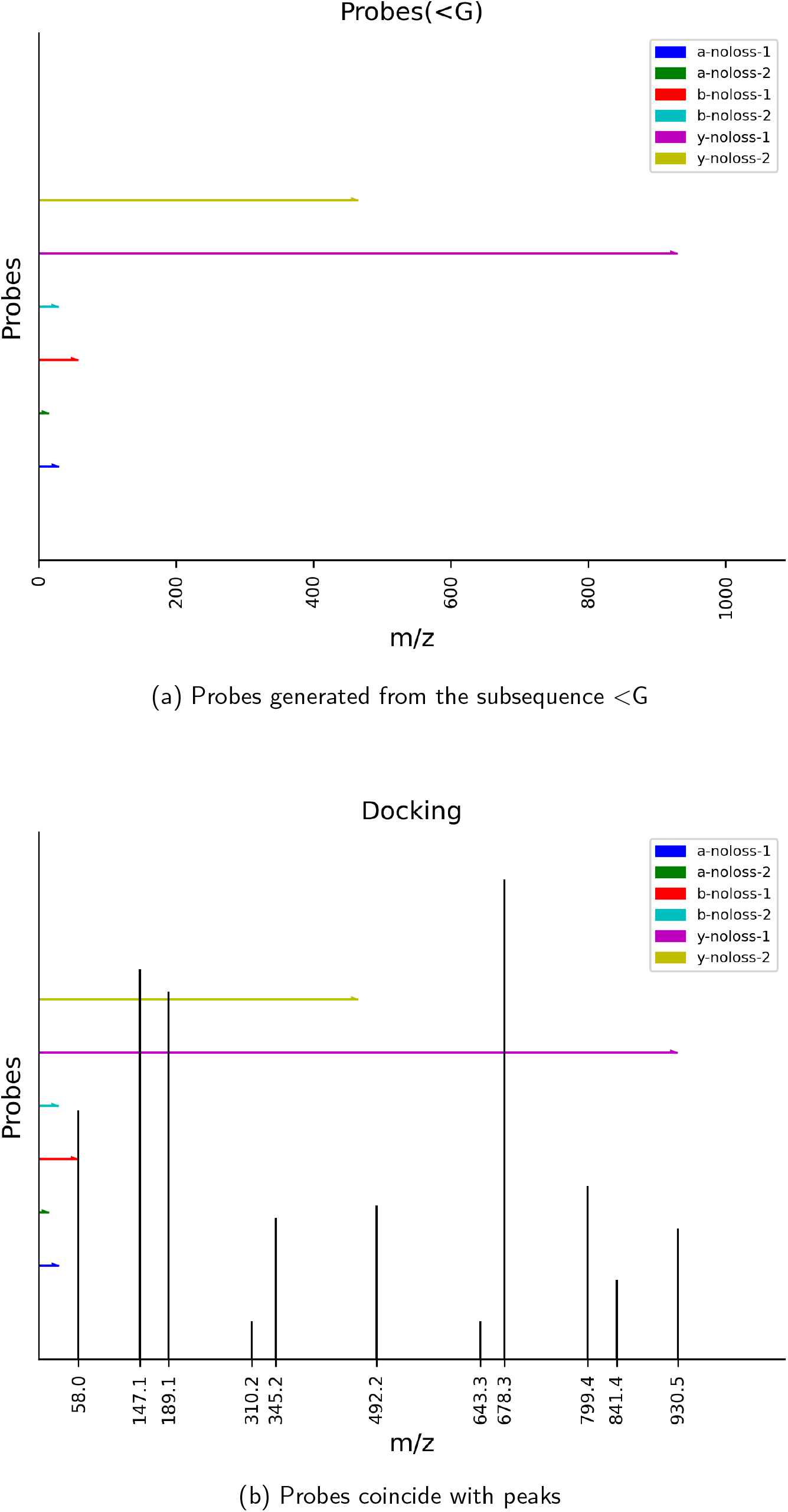
Probes and Docking. Probes are generated by simulated ionization, while peaks are obtained from a real mass spectrum. Docking probes and peaks helps determine if the assumed residue is the correct next residue in a peptide sequence.

Probe *j* is represented as *m*/*z*_*j*_ and the set of M probes is denoted as 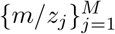. Spectrum probes are designed to identify which residue occupies the next position in a peptide sequence, requiring all candidate residues to be considered and converted into probes.

#### Docking

Theoretically, when a probe coincides with a peak (i.e., they share the same *m*/*z* value), it strongly suggests that both the probe and the peak originate from the same fragment ion. Consequently, the set of probes can confirm the existence of a corresponding real peptide subsequence. As illustrated in Figure 2b, the probe b-noloss-1 precisely hits the peak at *m*/*z* 58.0, indicating that the subsequence *<G* is indeed present in the peptide, thereby confirming glycine (*G*) as the next residue. Building on the subsequence <*G*, we continue to hypothesize the identity of the next residue (e.g., <*GM*, <*GF*, <*GA*, …) and construct probes for the extended 1-residue longer subsequence. By repeating this process, we can iteratively detect and confirm each subsequent residue, one at a time, until the precursor mass is exhausted.

We define **docking** as the process of comparing all peaks with all probes by calculating the differences in their *m*/*z* values. This comparison is used to infer whether the set of probes represents a real peptide subsequence. A **docking value** is defined as:

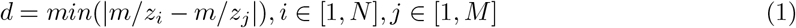

According to this formula, a smaller docking value increases the likelihood that the set of probes under investigation represents a genuine peptide subsequence. A docking value of zero is ideal, as it strongly indicates the true existence of the peptide subsequence.

#### Multi-layer Probes and Docking

Since all probes are compared with all peaks, the process of docking results in a large number of possible combinations. Occasionally, a docking value generated by an incorrect residue may happen to be smaller than that of the correct residue. In such cases, the wrong residue might be mistakenly selected as the next residue. To address this issue, we employ **Multi-layer Probes** to detect a chain of multiple subsequent residues, which enables us to obtain a more reliable docking value. The rationale behind this approach is that while an error may occur once, it is unlikely to happen repeatedly in succession. Theoretically, the more layers we consider, the more confident we become in selecting the correct next residue. Similar ideas have been successfully applied in other fields^41^, demonstrating their effectiveness.

As shown in Figure 3, the 1-layer probe supports that the first residue is a *G*, while the 2-layer probe supports that the first two residues are *G* and *M*. These results reinforce the conclusion that the first residue is *G*, enabling us to make the assertion confidently. As mentioned earlier, all candidate residues must be considered when generating probes. Consequently, if we have 20 residues, we need to generate 20 probe sets for 1-layer setting, and a total of 420 sets for 2-layer setting (20 + 20^2^ = 420). Let the number of layers be *L*. Then, the total number of probe sets that need to be generated is given by 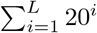, indicating that a significant amount of computation is required when calculating multi-layer probes, especially when *L* is large. To efficiently handle this computational load, we employed GPUs and CuPy^42^, a GPU-accelerated computing library for Python. The docking value *d* for multi-layer probes is calculated by averaging the docking values *d*_*i*_ of each layer:

**Figure 3.**
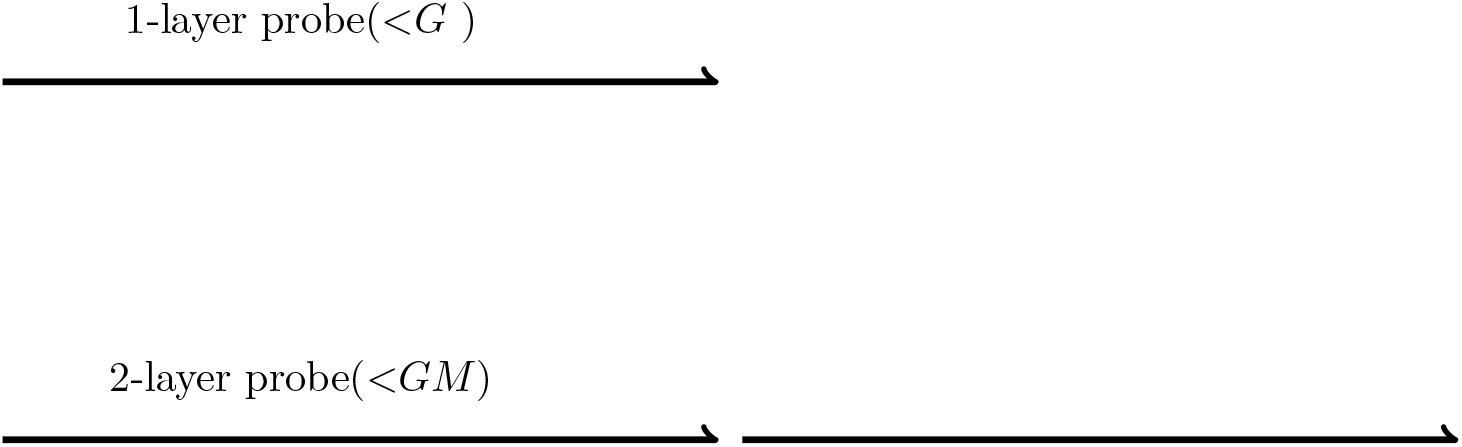
Probe and Multi-layer Probes. The 1-layer probe (top) detected <*G*, and the 2-layer probe (bottom) detected <*GM*.

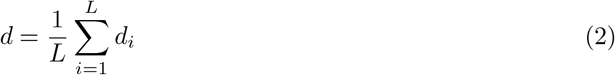

### 2.4 Dual Transformer model

Transformer^43^ is a deep learning model architecture introduced by Vaswani et al. in 2017. Unlike RNNs or CNNs, Transformer uses an attention mechanism to capture both short-range and long-range dependencies. This makes it particularly well-suited for our task because *de novo* peptide sequencing involves modeling relationships between peaks in a spectrum, which may be either closely spaced or far apart. In PepGo, we implemented a dual Transformer model to compute the probabilities of the next residues from two opposite ends of the peptide sequence (N-terminal and C-terminal), as shown in Figure 4. Since the N-terminal and C-terminal Transformer models share the same architecture, differing only in that they are trained on forward and reverse peptide sequences respectively, the following discussions apply to both models unless otherwise specified.

**Figure 4.**
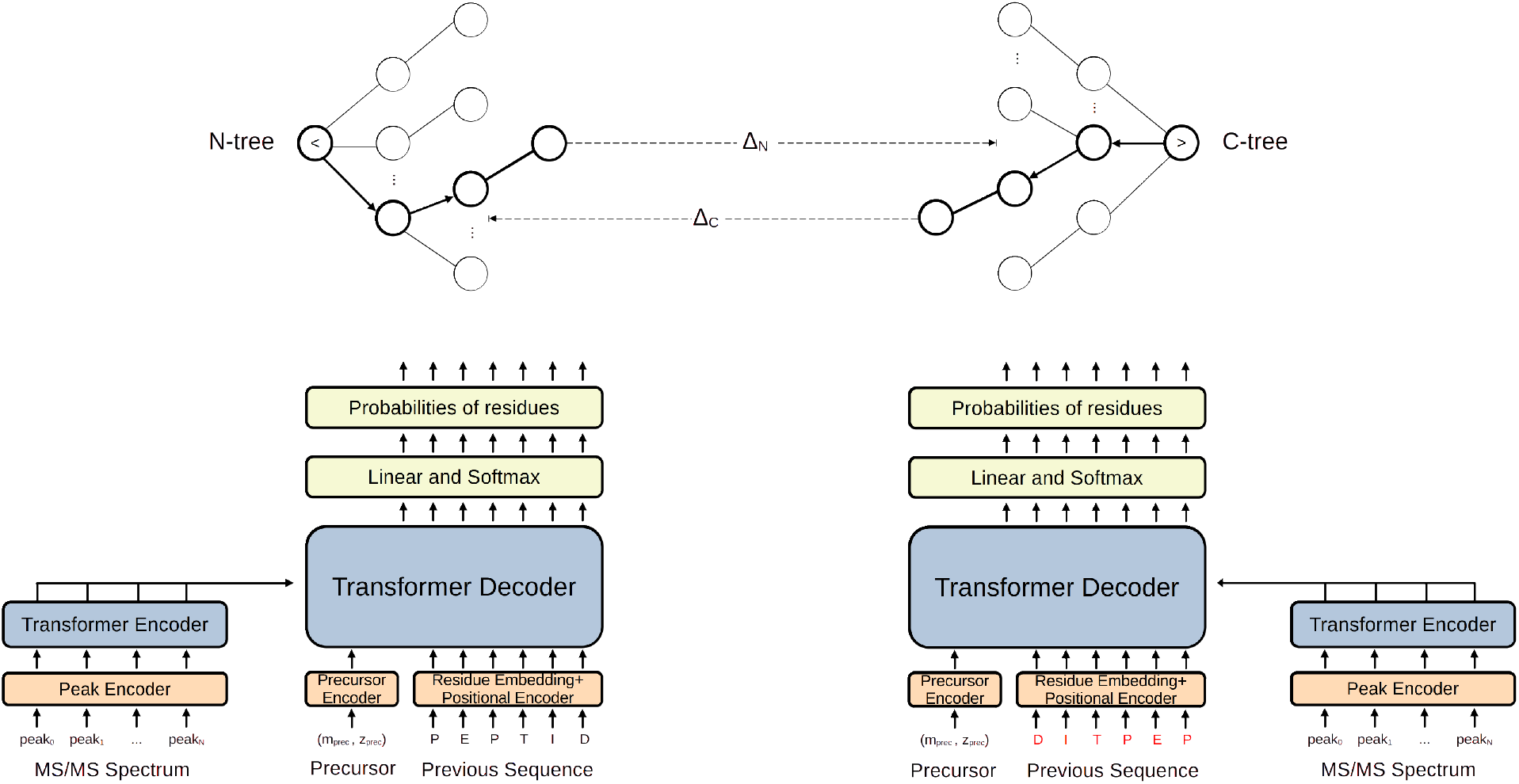
Overview of the PepGo model structure. (Bottom left) The N-terminal Transformer model for computing the probabilities of the next residues starting from N-terminal. This model’s decoder is trained on forward peptide sequences. **(Bottom right)** The C-terminal Transformer model for computing the probabilities of the next residues starting from C-terminal. Note that the ‘previous sequence’ is written as DITPEP to emphasize that the decoder is trained on the reversed version of the same peptide sequences. **(Top left)** The N-tree is constructed with *<* as the root node. The probabilities of the nodes in this tree are computed by the N-terminal Transformer model. **(Top right)** The C-tree is constructed with *>* as the root node. The probabilities of the nodes in this tree are computed by the C-terminal Transformer model. **(Top middle)** Δ_*N*_ and Δ_*C*_ are the rewards for the last nodes of N-tree and C-tree respectively.

We employed the Transformer neural network architecture from Casanovo^6^ as the foundational building block for our dual Transformer model, adapting its design to meet our specific requirements. We primarily discuss the differences in detail, while the components not specifically mentioned remain consistent with the original model.

#### Spectrum preprocessing

Unlike Casanovo, we do not discard peaks that fall outside a certain *m*/*z* range or lie close to the observed precursor mass. While such peaks may initially appear anomalous, they may arise from normal fragmentation processes and still provide crucial information for *de novo* peptide sequencing. For instance, high-charge ions, though possessing low *m*/*z* values, often carry important sequence-specific information. Rather than filtering peaks by their relative intensities, we retain all peaks under the assumption that low-intensity signals are not necessarily noise. Although noise is an inherent challenge in mass spectrometry data, our approach pins hope on the noise tolerance potential^44^ of Transformer models, especially when trained on large and diverse datasets. This minimizes the risk of discarding potentially valuable peaks prematurely. We also believe that filtering strategies should be developed with more sophisticated, context-aware methods that address noise at a deeper level. Additionally, we do not impose a hard upper limit on the number of peaks, as modern Transformer models can handle increasingly longer contexts^45, 46^, allowing for the efficient processing and analysis of spectra with a large number of peaks without sacrificing performance or accuracy.

#### Dual model structure

PointNovo and Casanovo use a beam search decoding strategy to predict each subsequent residue from one terminal to the other, which is straightforward and easy to implement. However, this approach overlooks the possibility of simultaneous prediction from both ends, leading to cumulative errors as prediction progresses, ultimately reducing accuracy. To mitigate this problem, PepGo adopts a dual-model structure that incorporates information from both ends. In this structure, the N-terminal Transformer model is trained on forward peptide sequences to compute probabilities exclusively for the next residues starting from the N-terminal. While the C-terminal Transformer model, trained on reversed peptide sequences, computes probabilities solely from the C-terminal, as shown in Figure 4. Both models share the same architecture, parameters, and hyperparameters.

### 2.5 Monte Carlo Double-Root-Tree Search

In state-of-the-art models of *de novo* peptide sequencing, such as DeepNovo ^7^, PointNovo^9^, and Casanovo^6^, beam search is conventionally used during the decoding process. At each iteration, beam search evaluates a fixed number (*top k*) of candidate sequences, continuing this exploration until the optimal prediction is found. However, the performance of beam search is limited by its inherent drawbacks. First, it retains only the *top k* sequences based on their current likelihoods, meaning its exploration is constrained by the beam width. While some sequences with lower initial likelihoods may eventually lead to better predictions, beam search may prematurely prune these paths. As decoding progresses, the risk of missing optimal solutions increases unless the beam width is significantly expanded, which can become computationally expensive. Additionally, beam search typically operates sequentially in one direction, making it difficult to coordinate predictions from both the N-terminal and C-terminal of a peptide. This sequential nature limits its ability to fully utilize contextual information from both ends of the peptide.

Monte Carlo Tree Search (MCTS) is a powerful algorithm for making optimal decisions^47^ and has been successfully applied to various decision-making problems, such as the game of GO ^11, 48^. Unlike beam search, which prunes paths early based on a fixed limit, MCTS explores a wider range of potential solutions by balancing exploration and exploitation over time. This allows MCTS to avoid prematurely discarding potentially optimal paths, thereby reducing the risk of missing optimal solutions. Additionally, the tree structure in MCTS enables the accumulation of knowledge about the decision space, refining future choices and making it especially powerful for long-term decisionmaking tasks. MCTS can efficiently handle much larger search spaces by strategically sampling and updating its tree structure, making it particularly well-suited for complex tasks. Moreover, MCTS determines one residue at each iteration, enabling predictions to alternate between the N-terminal and C-terminal. This bidirectional approach allows the peptide to be constructed from both ends concurrently, rather than from only one direction, as is the case with beam search.

The process of *de novo* peptide sequencing, which involves selecting the most appropriate next residue at each iteration, can similarly be framed as a decision-making problem for which MCTS is well-suited. However, to the best of our knowledge, MCTS has not yet been successfully applied in this domain. As an initial exploration, we propose and implement **Monte Carlo Double-Root-Tree Search (MCTTS)**, a variant of MCTS, to guide the decision-making process in *de novo* peptide sequencing. In this section, we provide a detailed explanation of the MCTTS algorithm. Since MCTS was originally developed for games^49^ and has been extensively used in that domain, many of its core definitions and concepts are rooted in game terminology. To better suit the requirements of our task, we have adapted these definitions and concepts from the standard MCTS algorithm^47^ and present them in a simplified manner.

MCTTS employs two search trees, each initialized with a different root node: the tree initialized with *<* is called **N-tree**, while the tree initialized with *>* is called **C-tree**, as shown in Figure 4. Each node *v* in the search trees represents a residue and has several properties: state *s*(*v*), reward *Q*(*v*) and visit count *N* (*v*). The **state** *s*(*v*) of a node contains the corresponding residue and its mass. For the root node, the residue is used to store the partially **determined residues** from previous iterations, while the mass stores the cumulative mass of the determined residues. Thus, the state of the root node acts as an accumulator, keeping track of residues that have already been determined and providing the context necessary for deciding on the next residue. The **reward** *Q*(*v*) of a node is a score that reflects the correctness of the corresponding residue. Notably, in canonical MCTS, the reward might be determined solely by simulation, whereas in MCTTS, it is derived from both internal simulations and external models, such as the probe model or Transformer model. The **visit count** *N* (*v*) of a node is a nonnegative integer that indicates the number of times the node has been visited so far. Each edge in the search trees connects a parent node to a child node, representing an action one residue may take to reach its subsequent residue, as shown in Figure 5.

**Figure 5.**
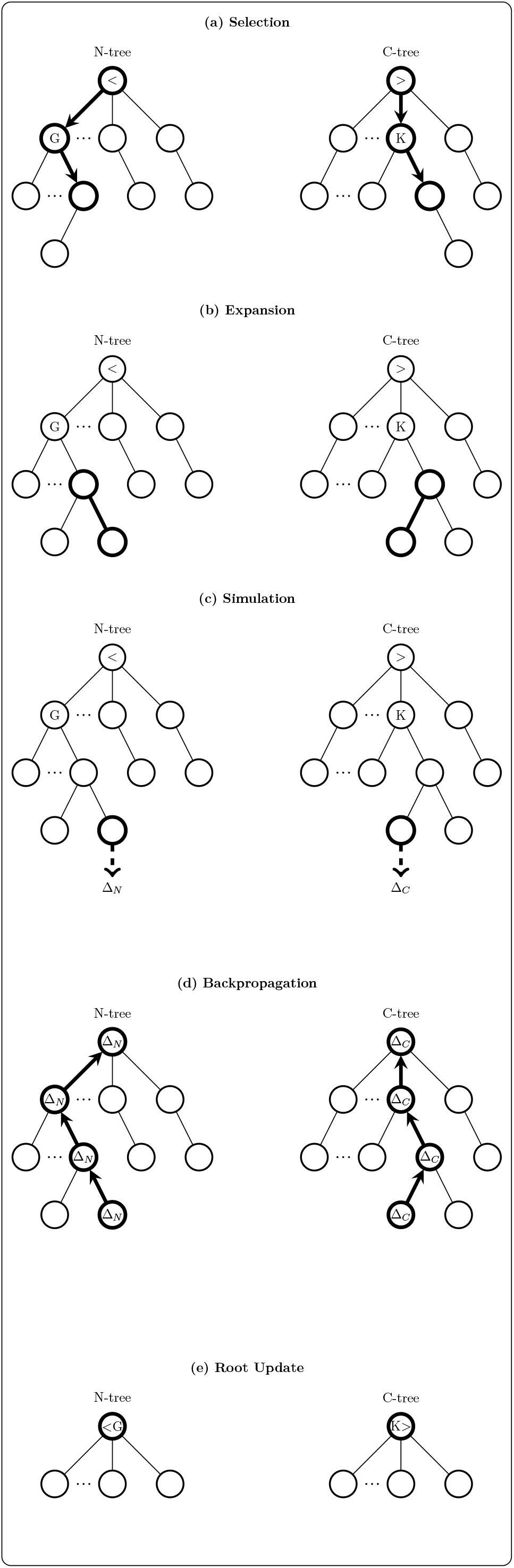
One iteration of the MCTTS algorithm consists of five steps. These steps are illustrated using an example peptide ‘GMRFWYK’. In the search trees, each edge represents an action connecting the current residue (represented by a node) to a subsequent residue. **(a)Selection:** In this step, the N-tree is initialized with *<* as the root. By recursively applying the tree policy, it first selects *G* as the best child and eventually reaches a leaf node. Similarly, the C-tree undergoes the same process, starting with *>* as the root, and initially selects *K* as the best child. **(b)Expansion:** One or more child nodes are added to the tree. **(c)Simulation:** From the terminal node added in the expansion step, a simulation is performed following the default policy, resulting in an outcome denoted as Δ (Δ_*N*_ for the N-tree and Δ_*C*_ for the C-tree). **(d)Backpropagation:** The outcome Δ is backpropagated through the selected nodes, updating their reward *Q*(*v*) and visit count *N* (*v*) sequentially. **(e)Root Update:** The best child of the root is determined as the next residue and appended to the determined residues. The status of the root is updated accordingly, and a new tree is then initialized with the updated root (*<G* for the N-tree and *K >* for the C-tree).

#### MCTTS consists of five steps in each iteration

##### Selection

Starting from the root node, a *tree policy* such as Upper Confidence Bound for Trees (UCT)^47^ is recursively applied to select the best child node until a leaf node is reached. Unlike the typical MCTS used in many games, the tree in MCTTS has a fixed depth, which can be configured and is primarily constrained by available memory and computational time.

##### Expansion

One or more child nodes are added to expand the tree, and the number of child nodes is up to the total number of candidate residues.

##### Simulation

A simulation is performed from the newly expanded node according to the *default policy*, and produces an outcome when it reaches the terminal state.

##### Backpropagation

The simulation outcome is backpropagated through the selected nodes, updating their statistics sequentially.

##### Root Update

The four steps above are repeated until the computational budget (a configurable number of attempts) is exhausted. Then, the best child of the root is selected and determined as the correct next residue, which is appended to the determined residues of the root. A new tree is then initialized with the updated root, preparing for the next iteration. After applying the same process to the other tree, the current iteration is completed.

Termination of iterations: N-tree and C-tree alternate in determining residues, with N-tree selecting a residue at the N-terminal, followed by C-tree choosing another residue at the C-terminal. This alternating approach ensures a balanced progression through the sequence. In this process, the precursor mass is a crucial constraint. Each time a residue is determined by either tree, its mass is subtracted from the precursor mass, and the resulting value is checked to see if it is equal to or less than zero. Ideally, the iterations should stop when the precursor mass is fully exhausted. However, unavoidable small mass errors can interfere with proper termination. For example, towards the end, the remaining precursor mass often does not reach exactly zero but instead may become a small positive value close to zero. This may cause the algorithm to initiate a new, unnecessary iteration and add an extra residue, resulting in incorrect predictions. To address this issue, we use the **remaining mass (**m_re_) instead of the precursor mass as the constraint, which is calculated by subtracting a proton mass from the precursor mass:

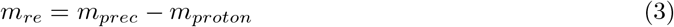

Since the smallest mass unit in a peptide is the mass of a hydrogen atom, referred to as the proton mass (*H*^+^), this mass is large enough to eliminate small mass errors while being small enough to avoid the opposite issue of predicting one residue fewer than the actual number.

Algorithm 1, written in pseudocode, illustrates the UCT algorithm applied in MCTTS. The algorithm begins by initializing a root node *v*_0_ with state *s*(*v*_0_), then the tree policy is applied to *v*_0_ and is recursively followed until the last node *v*_*l*_ is reached. Δ represents the reward for the terminal state reached by applying the default policy from the state *s*(*v*_*l*_). In each iteration, the mass of the newly appended residue, *m*_*new*_, is subtracted from the remaining mass *m*_*re*_. The iteration continues until *m*_*re*_ is exhausted, at which point *v*_0_ has been updated using its best child. Finally, the determined residues in state *s*(*v*_0_) are returned as the overall result. The constant *C*_*p*_ *>* 0 is used to balance exploration which encourages the selection of unvisited children, and exploitation which favors children that have shown promising results. Let *v* be the current node and *v*^*′*^ a child node of *v*. The state of node *v*^*′*^, denoted as *s*(*v*^*′*^), is returned by the function *model*(*s*(*v*)). The context information for this calculation includes the residues from the root *v*_0_ to node *v*, along with the corresponding mass of these residues. The function *model* reads *s*(*v*) and infers this context information to compute a reward, which is then assigned to node *v*^*′*^. Two models are available for calculating this reward: probe model and dual Transformer model.

Probe model uses the context information in *s*(*v*) as the input and calculate the docking value *d*_*v*_ of the node *v* with Formula 2. According to the Central Limit Theorem, *d*_*v*_ can be seen as approximately following a normal distribution, so we calculate the reward with a Gaussian activation function:

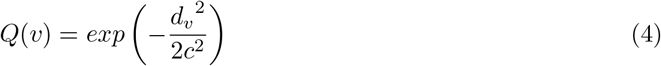

*c*=0.005 is the standard deviation, and the reward *Q*(*v*) ∈ (0, 1] can be regarded as a probability. Dual Transformer model outputs the probability of each node being the correct next residue, which is directly used as the reward *Q*(*v*).

Let **tail mass (**m_tail_) be the precursor mass minus the total mass of the selected residues in the current tree and the determined residues in the other tree, excluding *<* and *>*. Theoretically, if all these the residues are correct, the tail mass must precisely match the mass of a bag of residues. Such a bag of residues is actually a combination with repetition, and the number of such combinations is:

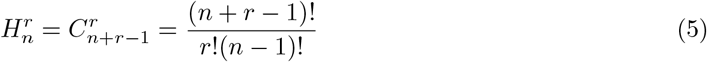

In Formula 5, *n* represents the number of different residues, and *r* is the maximum sequence length in consideration, which can be configured according to memory size. By enumerating all possible combinations, we get 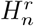 masses and sort them to obtain an array of ordered masses. We search the tail mass in this sorted mass array using binary search algorithm to find the closest mass *m*_*close*_, then calculate Δ with the function *g* below:

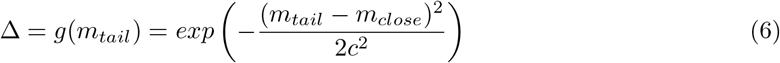

The constant *c* is the same as that in Formula 4, and Δ is the reward of playing out from the last node, which is the “Monte Carlo” part of our algorithm. Next, the reward value Δ is backpropagated along the path of the selected nodes, updating the statistics of each node. The visit count of each node is incremented by one, and its *Q*(*v*) value is updated by adding Δ. Finally, the root node *v*_0_ is updated by adding the residue of its best child to the determined residues.

An alternative method to calculate the reward value Δ is to perform a beam search from the last node until the tail mass is exhausted, then use the score of the best-fit peptide subsequence output by the Transformer model as the reward, referred to as “beam score”. This method maximizes the utilization of dual Transformer model and GPUs, while reducing the computational cost associated with binary search.

The combination of the Transformer model and the MCTTS algorithm represents a novel approach in *de novo* peptide sequencing, with MCTTS serving as a more powerful alternative to beam search. Similar methods have also been adopted across broader domains^50^.

The discussion above applies to both trees unless stated otherwise.

###### Algorithm 1

Upper Confidence Bound for Trees (UCT)

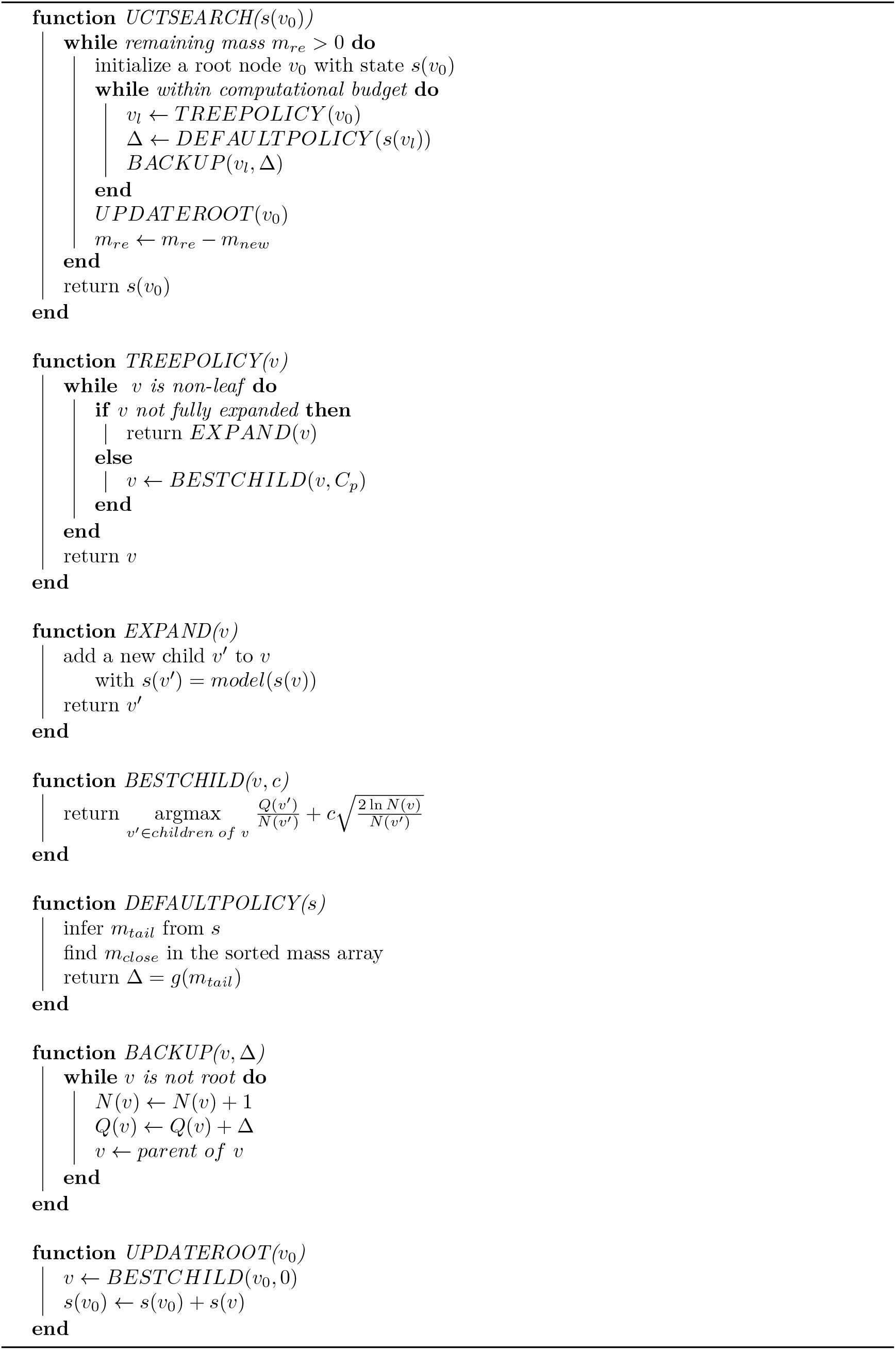

### 2.6 Quality Control

We employ the method proposed by Casanovo^6^ to filter out low-quality peptide predictions:

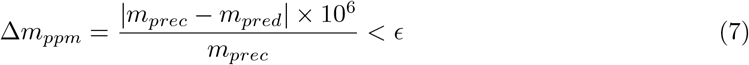

Δ*m*_*ppm*_ is the mass error in PPM (Parts Per Million), used to express a small difference between the precursor mass (*m*_*prec*_) and the total mass of the predicted peptide (*m*_*pred*_). The threshold value *ϵ* is configured based on either the properties of the mass spectrometer or prior knowledge and expertise. Peptide predictions that do not meet this threshold are excluded from the final results.

## 3. Experiments

To evaluate the performance of PepGo in *de novo* peptide sequencing, we conducted a comparative analysis against other state-of-the-art models, including Casanovo^6^, PointNovo^9^, and Novor^21^.The evaluation was conducted using synthetic peptides and their corresponding mass spectra, establishing a reliable and consistent environment to assess each model.

### Data Set

Most *de novo* peptide sequencing models, such as PointNovo and Casanovo, are trained using peptide-spectrum matches (PSMs)^6, 9^. Although the spectra are generated from mass spectrometry experiments, the actual peptides that generate these spectra are initially unknown, requiring database searches to identify the underlying peptides before training. As a result, PSMs are inherently error-prone and not fully reliable, even when filtered with high-quality standards. Training on these ambiguous PSMs biases the model, making it prone to producing wrong predictions.

Synthetic peptides, along with their generated spectra, offer several distinct advantages over PSMs for training *de novo* peptide sequencing models. First, synthetic peptides have known, chemically defined sequences, eliminating the risk of misassigned sequences or false positives commonly found in PSMs. This ensures that the training labels are always accurate and reliable. Additionally, the mass spectra generated from synthetic peptides are produced under controlled experimental conditions, resulting in clean, high-resolution spectra with fewer artifacts and less noise. Moreover, synthetic peptides can be engineered to include specific PTMs, enabling the tailored generation of spectra with known PTMs. Finally, synthetic peptides can be designed to systematically explore a wide range of peptide characteristics, such as varying lengths, amino acid compositions, charge states, and complex modifications.

We use synthetic peptides with their generated spectra^51–53^ as datasets for the training, validation, and testing of PepGo, as well as ground truth datasets to evaluate and compare the performance of different *de novo* peptide sequencing models. The synthetic peptide datasets in MGF format files were downloaded from **ProteomeTools** and converted to our custom .spec format. For duplicate peptides, one representative was randomly selected, and the others were discarded. Ultimately, we obtained a dataset consisting of 393,001 peptides with their corresponding spectra, which were divided into training(320,000), validation(40,000) and testing(33,001) sets. The datasets include two types of PTMs: oxidation on methionine (M)and carbamidomethyl on cysteine (C). Additionally, residues isoleucine (I) and leucine (L) were converted to X, as they have identical masses and cannot be distinguished currently.

### Model Training

We trained PepGo, Casanovo, and PointNovo on a server equipped with a single NVIDIA GTX2080 GPU.

### Parameters

PepGo operates in three distinct modes: probe, Transformer, and beam. In probe mode, PepGo utilized three types of fragment ions, *a, b* and *y*, with no consideration of neutral losses. The maximum charge state of the fragments was set to match the charge state of the precursor ion. Additionally, 2-layer probe was employed to enhance the probing mechanism. The reward *Q*(*v*) of nodes is calculated by docking value and Formula 4, while Δ is determined by Formula 6 based on tail mass. Notably, in this mode, PepGo can operate without training, as the docking value is derived from simple mathematical calculations rather than a trained model. In Transformer mode of PepGo, the configuration of dual Transformer model was aligned with that of Casanovo, allowing for a more direct comparison between the two models. *Q*(*v*) is produced by the Transformer model, while Δ is calculated using Formula 6 based on tail mass. In beam mode, the beam score is used as Δ, while *Q*(*v*) is produced by the Transformer model, as in Transformer mode.

In our implementation of MCTTS, the search tree was constructed with a depth of 3, and the computational budget was limited to 20 iterations, ensuring a balance between computational efficiency and search accuracy. When calculating Δ value using the function *g*, the length of peptide sequence under consideration was set to 13 residues.

For Casanovo, the number of peaks was set to 300, with the batch size for training and prediction set to 64 and 256, respectively. All other parameters were kept as default. For PointNovo, the batch size for training was limited to 8 due to the memory constraints of the RTX 2080 GPU, and the number of epochs was set to 10, as the optimal model was typically achieved early in the training process. All other parameters were consistent with the default settings. For Novor, we used the publicly available, pre-trained version for comparison.

### Metrics

We use peptide recall and residue recall as the metrics to evaluate all the models. **Peptide Recall** is defined as the number of correctly predicted peptides divided by the total number of ground truth peptides. It is the most stringent criterion, as it requires exact matches between the predicted and ground truth peptide sequences, where even a single mismatched residue results in a failed match. Therefore, peptide recall is a valuable metric for evaluating a model’s real-world performance in practical applications. **Residue Recall**, on the other hand, is defined as the number of correctly predicted residues at the same position as in the ground truth peptide sequences, divided by the total number of residues in those sequences. While less stringent than peptide recall, residue recall still provides a useful measure of an model’s overall accuracy.

## Results

As shown in Figure 6a, PepGo achieved the highest peptide recall at 71.6%, outperforming Casanovo(64.5%), PointNovo(58.8%), and Novor(37.1%). A similar trend is observed in the comparison of residue recall presented in Figure 6b. The Venn diagram in Figure 6c illustrates the peptide identification overlap among the four models. A total of 6,913 peptides were identified by all the models, showcasing a significant shared capability in peptide prediction. PepGo identified 1,887 unique peptides that were not predicted by any of the other models. Notably, although PepGo and Casanovo share 19,773 peptides, PepGo identified 3,870 unique peptides that Casanovo was unable to detect, as shown in Supplementary Figure S2a. This indicates that MCTTS functions quite differently from beam search, enabling it to explore the peptide space in a distinct manner. We also examined peptide recall across different peptide lengths, as depicted in Figure 6d. All approaches displayed a consistent trend: recall decreases as length increases. This decline may be attributed to error accumulation as peptide sequence length increases, resulting in progressively unreliable predictions.

**Figure 6.**
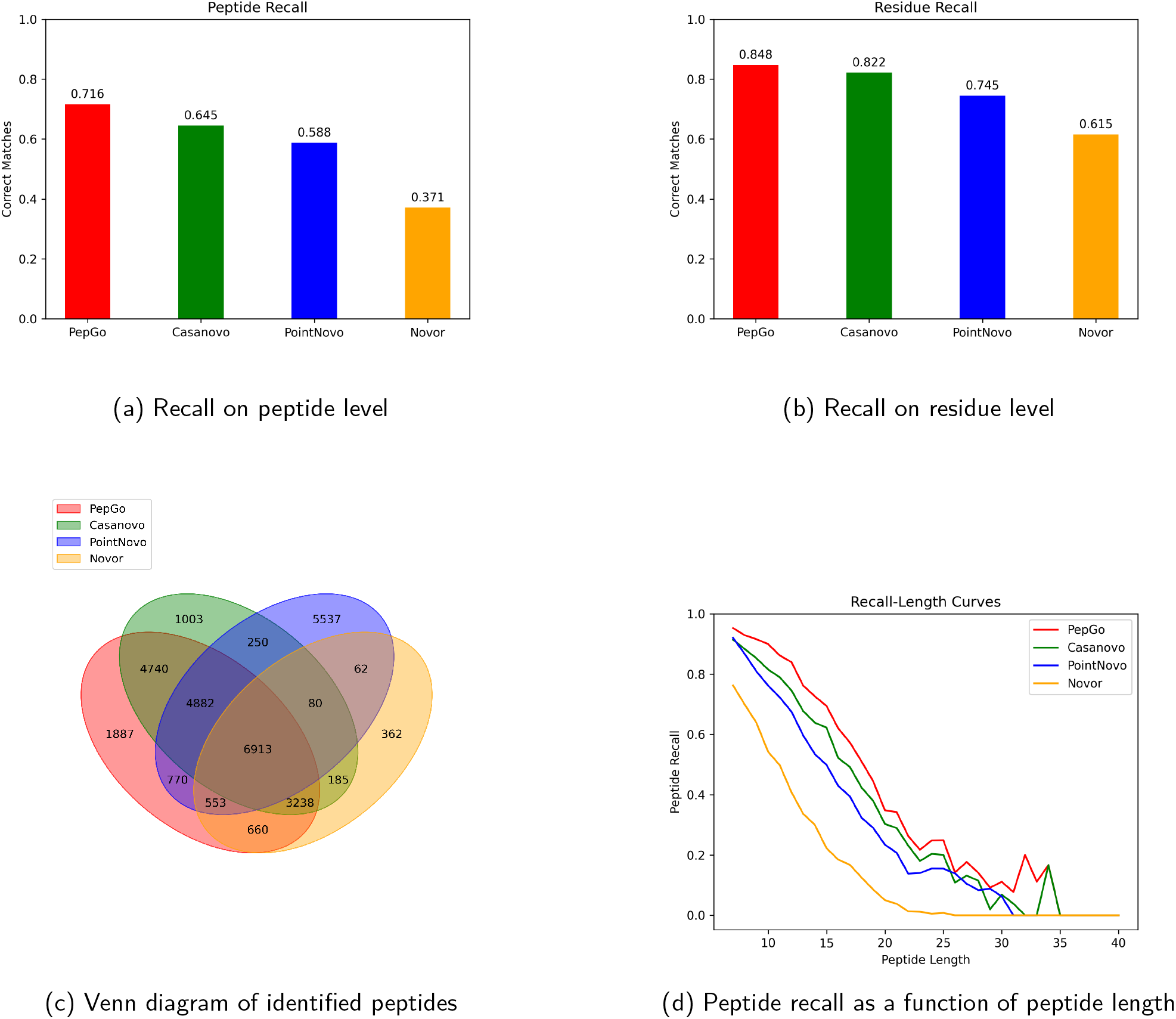
Comparison of PepGo’s performance with Casanovo, PointNovo, and Novor. **(a)** PepGo achieved the highest peptide recall among all the models. **(b)** PepGo achieved the highest residue recall. **(c)** The venn diagram shows that 6,913 peptides were identified by all four models, while PepGo uniquely identified 1,887 peptides that were not found by any of the other models. **(d)** All models display a decline in peptide recall as peptide length increases, with PepGo leading in overall performance, especially for short to mid-length peptides.

We also compared the performance of PepGo’s three modes with the other three models. As shown in Supplementary Figure S1, PepGo operating in Transformer mode(*PepGo*_*T*_) achieved the highest peptide recall(70.1%) and residue recall(84.3%), demonstrating the superior performance of the combination of Transformer model and MCTTS.

## 4. Discussion

*De novo* peptide sequencing is a challenging yet fundamental problem in science. To address this problem, we developed PepGo by combining Transformer model with Monte Carlo Double-Root-Tree Search (MCTTS). The self-attention mechanism in the Transformer model allows PepGo to capture long-range dependencies between different peaks in a spectrum and residues in a peptide sequence, thereby enhancing its ability to compute the probabilities of the next residue. MCTTS provides PepGo with a broader search scope compared to beam search by considering all candidate residues, rather than limiting the search to the *top k* residues. PepGo can operate in probe mode, where it can make predictions without training. While this mode might have varying prediction accuracy, it is particularly useful when the required training data, such as peptides with rare PTMs, is unavailable. We trained PepGo using synthetic peptides and their corresponding spectra. These synthetic data provide a more accurate and reliable training set, which can be designed in advance to meet specific requirements. PepGo is a novel approach and it has demonstrated superior performance in our experiments.

## 5. Author Contributions

Yuqi Chang designed the model and algorithm, implemented the code, conducted experiments, analyzed data, and drafted the manuscript. ChatGPT was utilized solely for checking grammar, without contributing to the scientific content. Siqi Liu and Karsten Kristiansen co-supervised the work and provided revisions to the manuscript.

## 6. Code availability

The source code of PepGo is in this github repo: https://github.com/alifare/PepGo

## A Supplementary Figures

**Figure S1:**
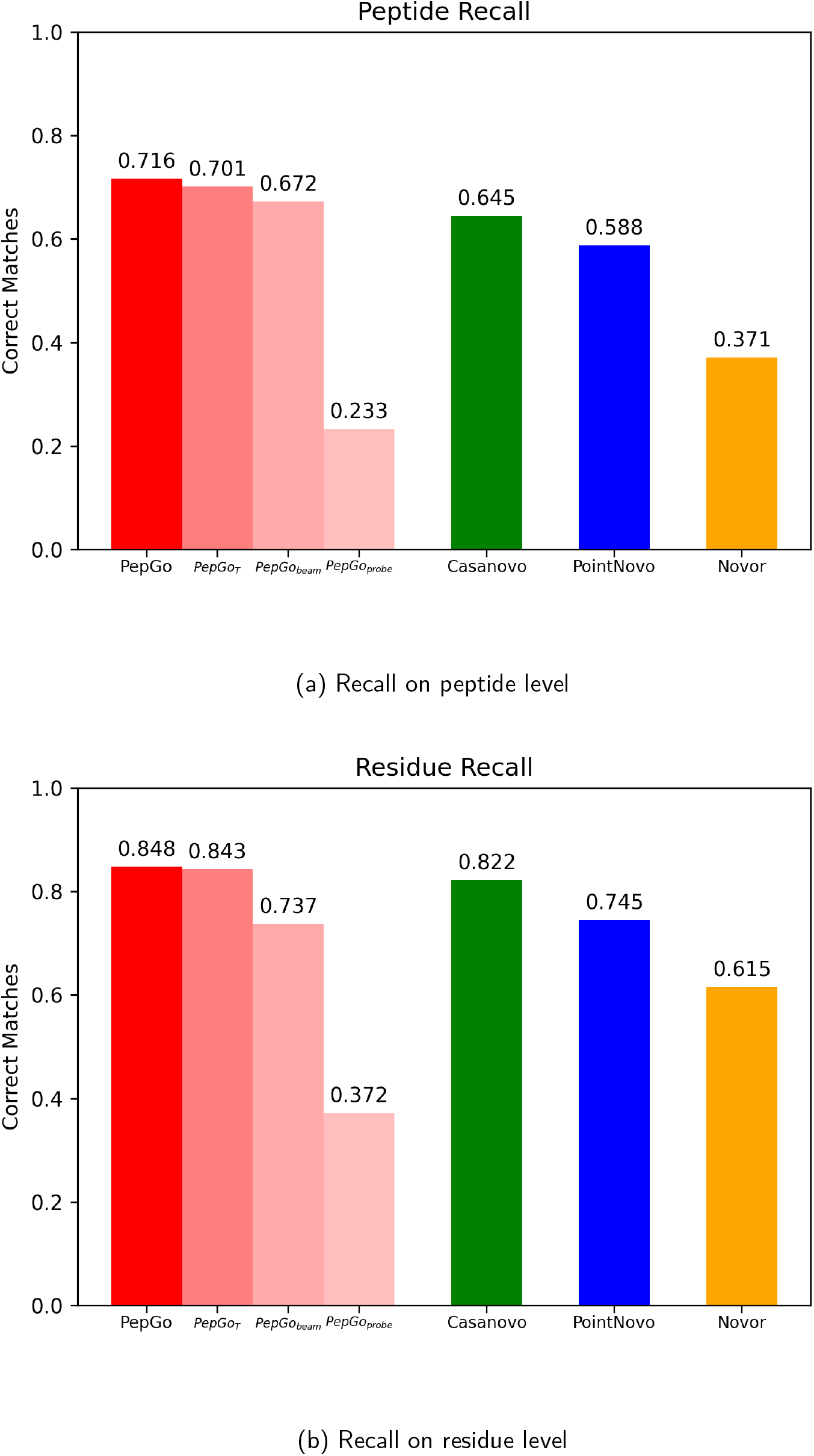
Comparison of the performance of PepGo’s three modes with Casanovo, PointNovo, and Novor. *PepGo*_*probe*_, *PepGo*_*T*_ and *PepGo*_*beam*_ represent the results of PepGo working in probe mode, Transformer mode and beam mode, respectively. PepGo represents the overall result, generated by replacing incorrect predictions from *PepGo*_*T*_ with the corresponding correct predictions(if available) from the other two modes.

**Figure S2:**
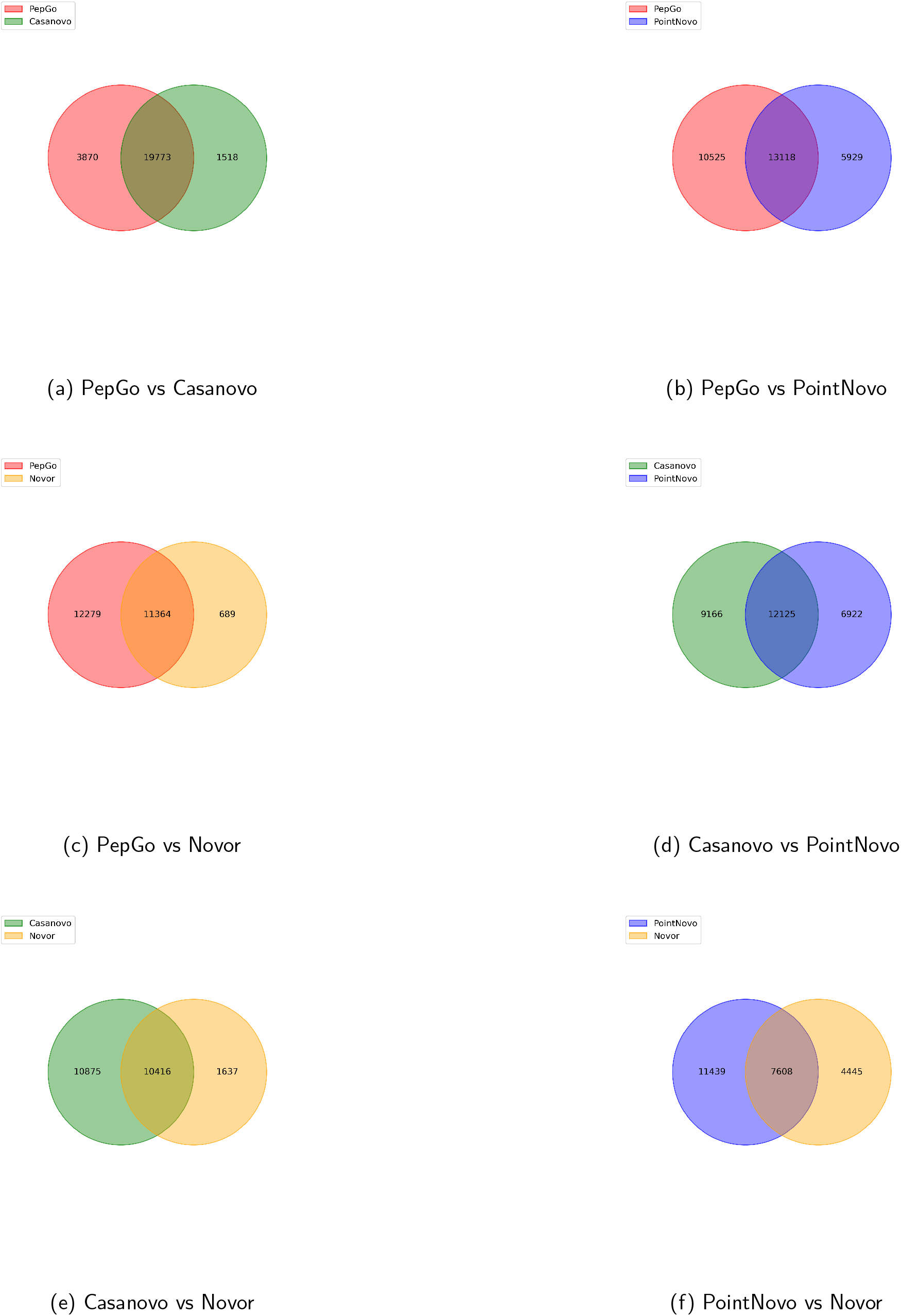
Two-set Venn diagrams illustrating the overlap of peptides identified by PepGo, Casanovo, PointNovo, and Novor. Each diagram compares two models, showcasing the shared peptides identified by both, as well as the unique peptides specific to each model, providing insight into their comparative performance in peptide identification.

